# Marine predator spatial conservation priorities are taxon-specific

**DOI:** 10.1101/2023.03.02.530743

**Authors:** Elizabeth Boyse, Simon J. Goodman, Maria Beger

## Abstract

Marine predators are globally threatened by anthropogenic stressors, but are key for ecosystem functioning. Their worsening conservation statuses indicate that current management is failing, requiring us to urgently reimagine their conservation needs to ensure their survival. Their life histories, threats, and resource needs are diverse. Consequently, spatial conservation areas targeting all species will overlook such heterogeneity, contributing to the problem. Here, we demonstrate that marine mammals, elasmobranchs and teleost fishes return drastically different spatial conservation priority areas, based on Marxan scenarios for 42 marine predator species in the Mediterranean Sea. None of the marine predators are sufficiently covered by the current marine protected area (MPA) system, with marine mammals being the least protected despite having the greatest designated MPA extent, highlighting disconnects between conservation goals and current management outcomes. To save marine predators, taxon specific ecological requirements and resulting spatial heterogeneity need to be accounted for in marine spatial planning.

## Introduction

Marine predators (mammals, sharks, rays and bony fishes) are globally declining due to anthropogenic pressures including fishing, climate change and habitat degradation (Avila et al., 2018; Dulvy et al., 2021). Their loss reduces important ecosystem functions such as top-down control, redistribution of nutrients, habitat engineering, and carbon sequestration (Hammerschlag et al., 2019). Marine predators are typically either protected within marine protected areas (MPAs) planned across multiple species and habitats, or focused on popular taxa, *i.e*., marine mammals (Notarbartolo di Sciara et al., 2016), but both approaches neglect the heterogeneous habitat requirements of predators stemming from their complex life histories, contributing towards their ineffectiveness (Klein et al., 2015). Further, marine predator home ranges typically exceed plausible sizes for MPAs, allowing exposure to threats outside MPA boundaries (Conners et al., 2022).

Over a third of marine mammals and elasmobranchs are classified as threatened in IUCN Red List assessments, and many commercially important predatory fish populations have already collapsed (MacKenzie et al., 2009; Avila et al., 2018; Dulvy et al., 2021). Fishing is the largest threat to all marine predators through resource exploitation, direct harvesting and bycatch. Further, climate-induced habitat degradation and range shifts are increasingly prominent threats (Avila et al., 2018; Dulvy et al., 2021). The impacts of shipping traffic are well established for marine mammals, but less certain for sharks or teleost fish (Schoeman et al., 2020). The loss of marine predators from ecosystems causes trophic cascades and reduced resilience to climate change (Estes et al., 2016). They also have high cultural and economic significance which can be both a benefit (*i.e*., high conservation interest) and a detriment (*i.e*., drives high demand) (Estes et al., 2016). Therefore, we urgently need to improve understanding of the conservation requirements for top marine predators to prevent further declines.

Marine spatial planning (Directive 2014/89/EU) offers a coordinated and transparent approach to managing different stakeholders using the marine space, whilst minimising impacts to the environment. Incorporating marine predators into spatial planning is important to indicate ecologically significant areas, *i.e*., with high productivity, species diversity or biomass of prey species (Augé et al., 2018). Incorporating multi-taxa approaches to identify priority areas has been advocated (Augé et al., 2018), but this oversimplifies the diverse habitat requirements of different taxonomic groups (Heupel et al., 2019). In contrast, predator-specific MPAs focus on narrower objectives, *i.e*., protecting specific life stages such as breeding or feeding grounds, but this strategy will only be effective if the protected life history stages maximise population growth rates (Conners et al., 2022). Different marine predator taxa have distinct spatial requirements, affecting their susceptibility and exposure to different threats (Avila et al., 2018). For example, divergent thermoregulatory strategies mean that marine mammals represent highest predator richness in temperate and polar waters, and in pelagic zones, while sharks and teleost fishes dominate tropical and coastal waters (Grady et al., 2019). At finer-scales, spatial partitioning between species is driven by mechanisms such as competitive exclusion and varying life history strategies (Heupel et al., 2019). Attempting to maximise conservation benefits for taxonomic groups offers a suitable balance between species-specific approaches, which will not afford protection to unstudied species, and broad (all taxa) biodiversity objectives that fail to account for taxon-specific requirements.

The Mediterranean Sea hosts a high diversity of marine predators that are exposed to some of the highest human impacts globally (Coll et al., 2010). Consequently, the Mediterranean Sea is an extinction risk hotspot for elasmobranchs (Dulvy et al., 2021), and local extinctions of marine mammal and teleost fish populations have already occurred (Bearzi et al., 2008; MacKenzie et al., 2009). Only 6 % of the Mediterranean Sea is covered by MPAs, and of these, 95 % have no regulations in place, owing to most being coastal and coinciding with high vessel density areas resulting in stakeholder conflicts (Claudet et al., 2020). Transboundary marine spatial planning has been encouraged given the large number of relatively small countries bordering the Mediterranean Sea (Li and Jay, 2020), yet marine spatial planning for marine predators has so far focused on single species or small spatial scales (Mazor et al., 2016; Carlucci et al., 2021). To ensure that spatial planning results in the best conservation benefits for marine predators, it needs to be executed at scales relevant to the expansive spatial ranges of marine predators (Estes et al., 2016; Conners et al., 2022).

In practice, conservation practitioners use all available biodiversity information for spatial planning prioritisations across taxa, and typically omit taxon-specific requirements. In this paper, we test whether taxa with different spatial ranges and habitat requirements need different conservation priority areas in the Mediterranean Sea. We firstly model the distributions of marine mammals, elasmobranchs, and large teleost fishes. Second, we identify separate and joint reserve prioritisation solutions for different taxa to test our expectation that each taxon requires specific conservation areas. Finally, we evaluate how different our reserve networks are compared to currently designated MPAs. We highlight discrepancies in the realised and required conservation efforts for marine predators and develop recommendations to facilitate the implementation of improved management measures for each group.

## Methods

### Species distribution modelling

We classify a marine predator as having a total length ≥100 cm and a trophic level ≥4 based on FishBase (https://www.fishbase.se/) or SeaLifeBase (https://www.sealifebase.ca/) records (Boyse et al., 2023). Setting a threshold of ≥40 occurrence records from GBIF (https://www.gbif.org, June 2020, GBIF Occurrence Download https://doi.org/10.15468/dd.tqx2he), OBIS (https://obis.org/), EurOBIS (https://www.eurobis.org/), the Mediterranean Large Elasmobranchs Monitoring (Medlem) database and Accobams (ACCOBAMS Survey Initiative, 2020; Mancusi et al., 2020), data were available to model distributions of 42 marine predator species, covering three taxonomic groups (20 teleost fishes, 13 elasmobranchs, 9 marine mammals). We obtained data for six environmental predictors (bathymetry, bathymetric slope, chlorophyll *a*, distance from shore, sea surface temperature mean and sea surface temperature range) from Bio-ORACLE and Marspec in WGS84 projection and 0.83° x 0.83° resolution (Sbrocco and Barber, 2013; Assis et al., 2018). Prior to modelling, we spatially thinned occurrence records with a nearest neighbour distance of 10 km (Aiello-Lammens et al., 2015). We modelled species distributions using maximum entropy (MAXENT), multiple adaptive regression splices (MARS) and random forest (RF) algorithms with the SSDM R package (Phillips et al., 2006; Schmitt et al., 2017). We generated 10, 000 background points for MAXENT. MARS and RF require pseudo-absence data, which we created randomly using the two-degree method, with 1000 points for MARS and an equal number of pseudo-absences as presence data for RF (Barbet-Massin et al., 2012). We made ensemble models across the different algorithms using weighted AUC scores. We converted ensemble habitat suitability models for each species to presence-absence models using the sensitivity equals specificity threshold (Schmitt et al., 2017). Binary ensemble models were summed to produce stacked species distribution models to visualise patterns in species richness.

### Priority areas for marine predators in the Mediterranean

We divided the Mediterranean Sea into planning units of 10 km x 10 km, in line with European Union guidelines on spatial planning (European Commission, 2007), resulting in 25,141 planning units in total, in the Lambert Azimuthal Equal Area projection (EPSG:3035). We assigned each planning unit an opportunity cost of displaced vessel traffic, represented by annual vessel density (hours per square kilometre) at a 1 km x 1 km resolution (European Marine Observation and Data Network, EMODnet; https://www.emodnet.eu/). We averaged annual vessel density across the available four years (2017-2020) and summed the data to a 10 km x 10 km resolution. Vessel density is a suitable surrogate for opportunity cost as this incorporates multiple sectors, including fishing, cargo, passenger and tankers which represent important threats for marine predators (Avila et al., 2018; Dulvy et al., 2021).

We employed the decision support software Marxan to identify priority areas for marine predator conservation. Marxan provides near-optimal solutions to the minimum set problem where conservation features (*i.e*., species) are adequately represented for the least possible cost (Ball et al., 2009). Our conservation features consisted of individual marine predator binary species distributions for 42 species. We set a target of 30 % protection for each species following the post-2020 Global Biodiversity Framework guidance (CBD, 2021). We ran Marxan using the simulated annealing algorithm and a boundary length modifier of 0.01 after calibration. We performed 100 iterations for each of the conservation feature scenarios: 1) all marine predator species, 2) marine mammals, 3) teleost fishes and 4) elasmobranchs.

### Comparing conservation planning scenarios with different taxonomic information

We used both the selection frequency, *i.e*., how many times each planning unit was selected in 100 iterations, and the ten solutions with the lowest objective scores to compare conservation priority differences across marine predator taxa. First, we quantified the overlap between planning unit selection frequencies of different taxa with Cohen’s Kappa coefficient (McHugh, 2012). The Kappa statistic requires categorical data so we classified the selection frequencies into five groups, 0, <25, 26-50, 51-75, >75 (Ruiz-Frau et al., 2015). Second, we performed hierarchical clustering with Jaccard dissimilarities from the 10 best solutions across the conservation feature scenarios (Brumm et al., 2021). We also compared the average cost and number of planning units required across the different taxa.

We downloaded the most current database for MPAs in the Mediterranean from MAPAMED, which includes 1,126 designated MPAs (MedPAN and SPA/RAC, 2022)(Figure 1B). We included MPAs with a national statute, Natura 2000 sites, and Specially Protected Areas of Mediterranean Importance (SPAMI) (MedPAN and SPA/RAC, 2022). We calculated the overlapping area between species distributions and MPAs and our ten best spatial prioritisation solutions to quantify which taxa are currently receiving most protection, and differences among taxa-specific prioritisation solutions.

**Figure 1.**
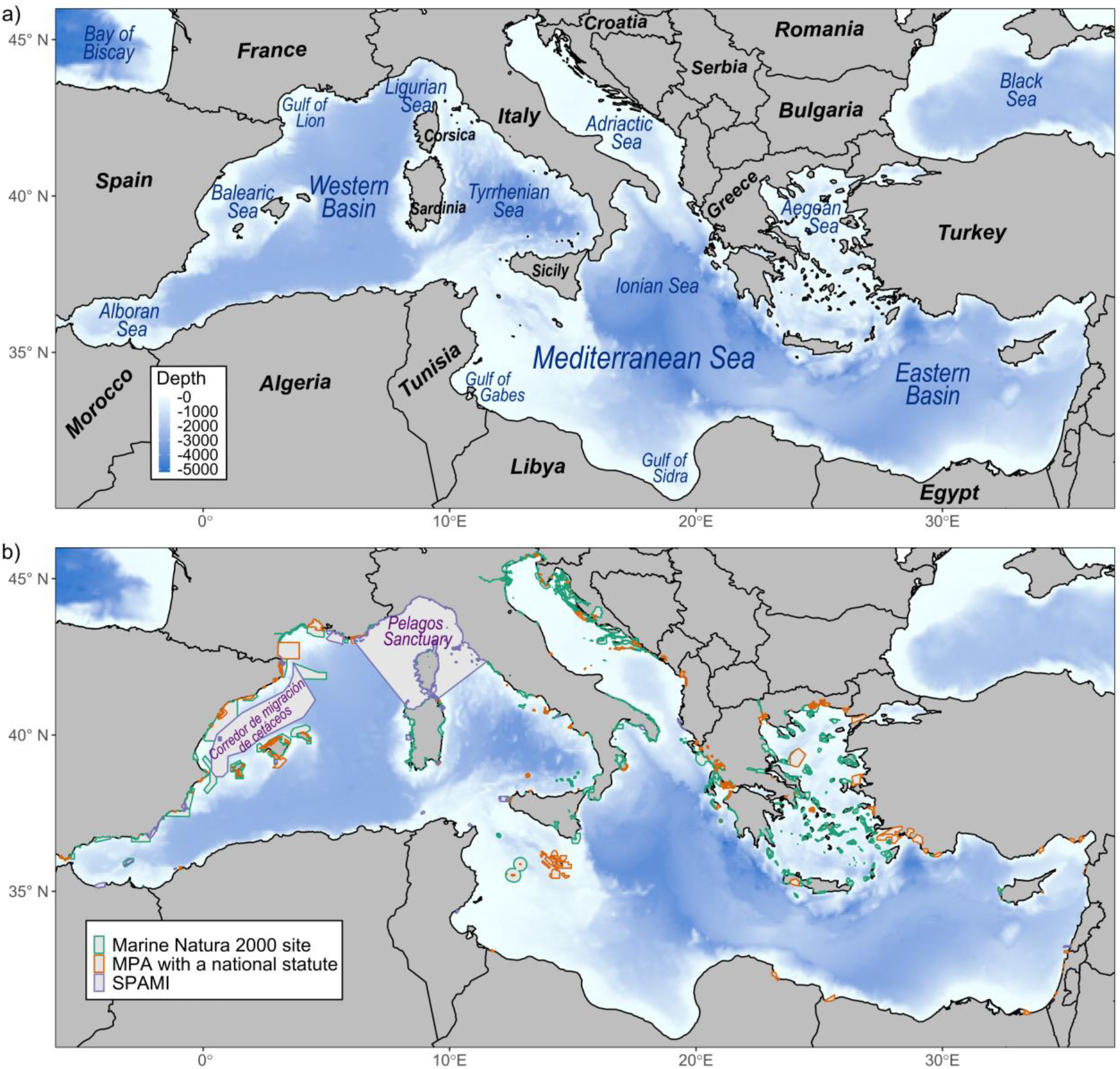
Bathymetric maps of the Mediterranean Sea showing a) marine regions, and b) designated marine protected areas including Marine Natura 2000 sites, marine protected areas (MPAs) with a national statute and Specially Protected Areas of Mediterranean Importance (SPAMIs).

## Results

Different taxa of marine predators have distinct distribution patterns (Figure 2). High species richness of elasmobranchs and teleost fishes is found along the coastlines of the north-western basin as well as the Balearic Islands, Corsica and Sardinia. Elasmobranchs have wider distributions in the Adriatic Sea, Aegean Sea and along the coastlines of Tunisia and Sicily, compared to teleosts. Highest species richness of marine mammals occurs in the Alboran Sea and between the Balearic Islands and Corsica/Sardinia. Overall, there is clear decrease in species richness with distance from shore, with highest species richness occurring in the north-western basin as well as the Adriatic and Aegean Seas.

**Figure 2.**
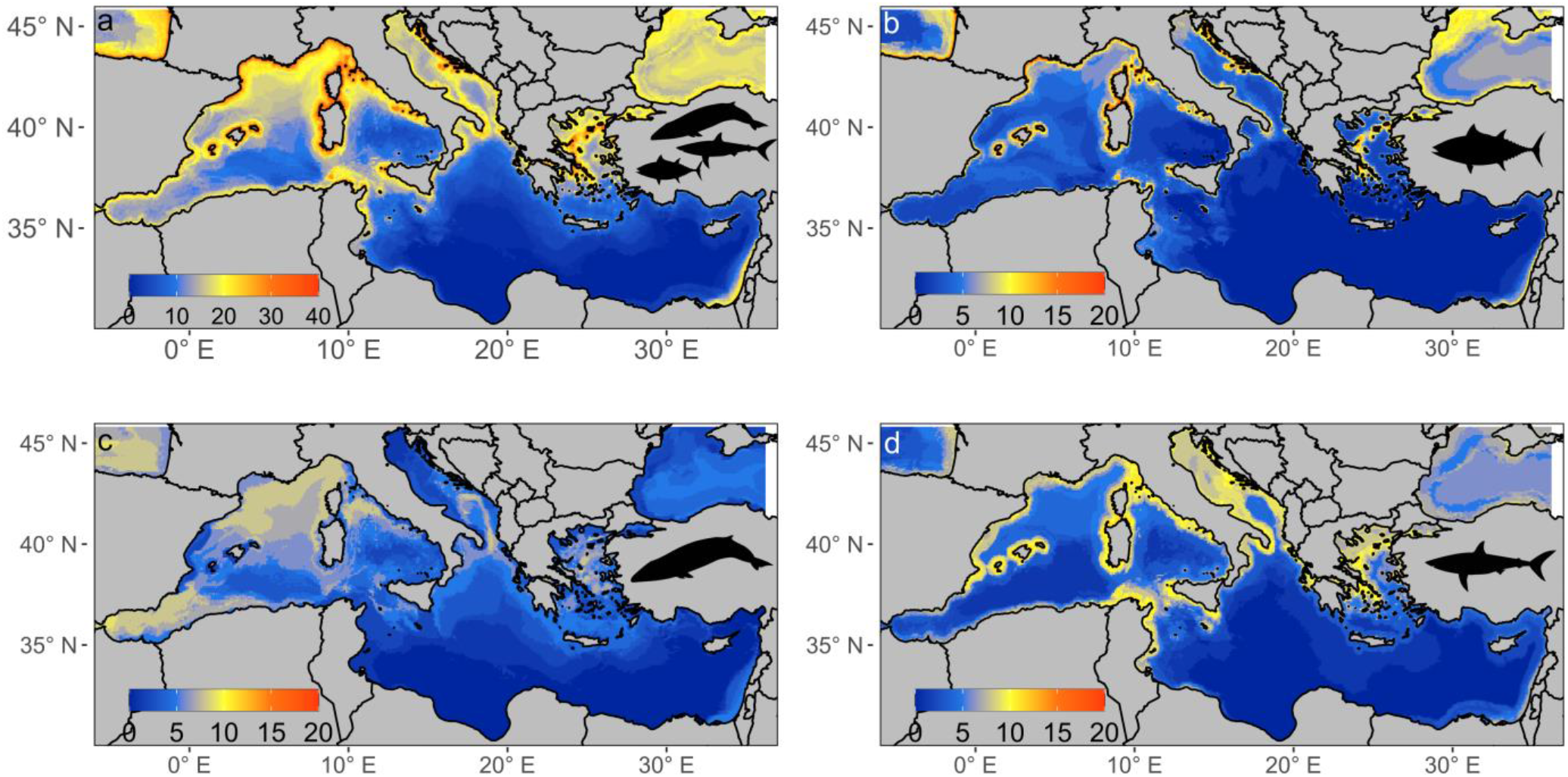
Maps of the Mediterranean Sea showing species richness from stacked species distribution models of a) all taxa, b) teleost fishes, c) marine mammals, and d) elasmobranchs.

Conservation feature scenarios considering taxa separately resulted in vastly different spatial prioritisation solutions (Figure 3). The Kappa statistic and hierarchical cluster analysis show highest similarity between selection frequencies for elasmobranchs and all taxa (Figures 4). Visually, elasmobranchs and all taxa scenarios share similar high selection frequency areas occurring along the coastlines of Tunisia and Egypt, the southern Alboran Sea, and the northern Aegean Sea. The Kappa statistic shows mammals and teleosts to have similar disagreement to the all taxa scenario, while cluster analysis reveals greatest dissimilarity between mammals and all taxa.

**Figure 3.**
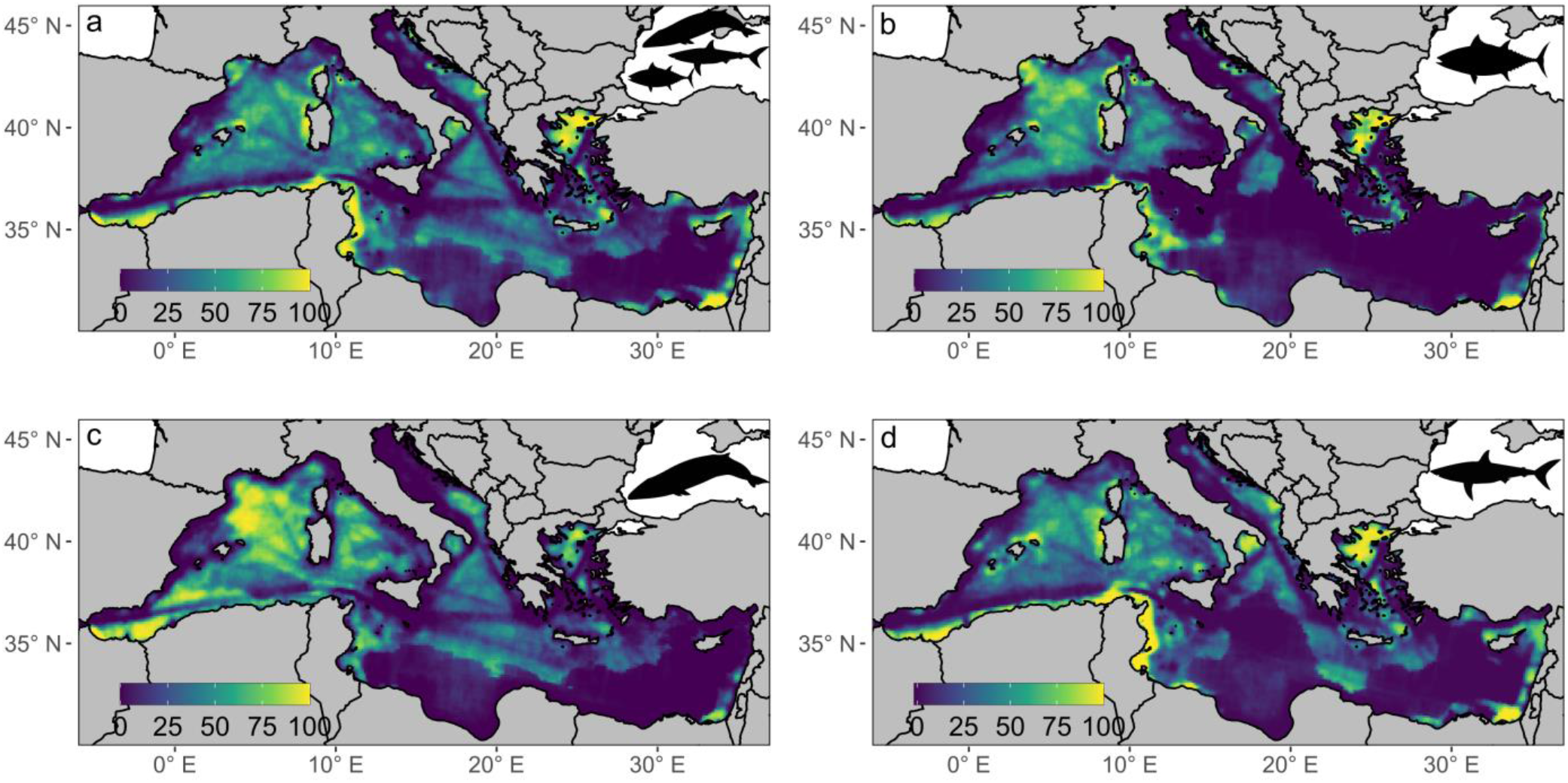
Maps of the Mediterranean Sea showing the selection frequency from Marxan outputs for each of the taxa scenarios; a) all species, b) teleost fishes, c) marine mammals, and d) elasmobranchs.

**Figure 4.**
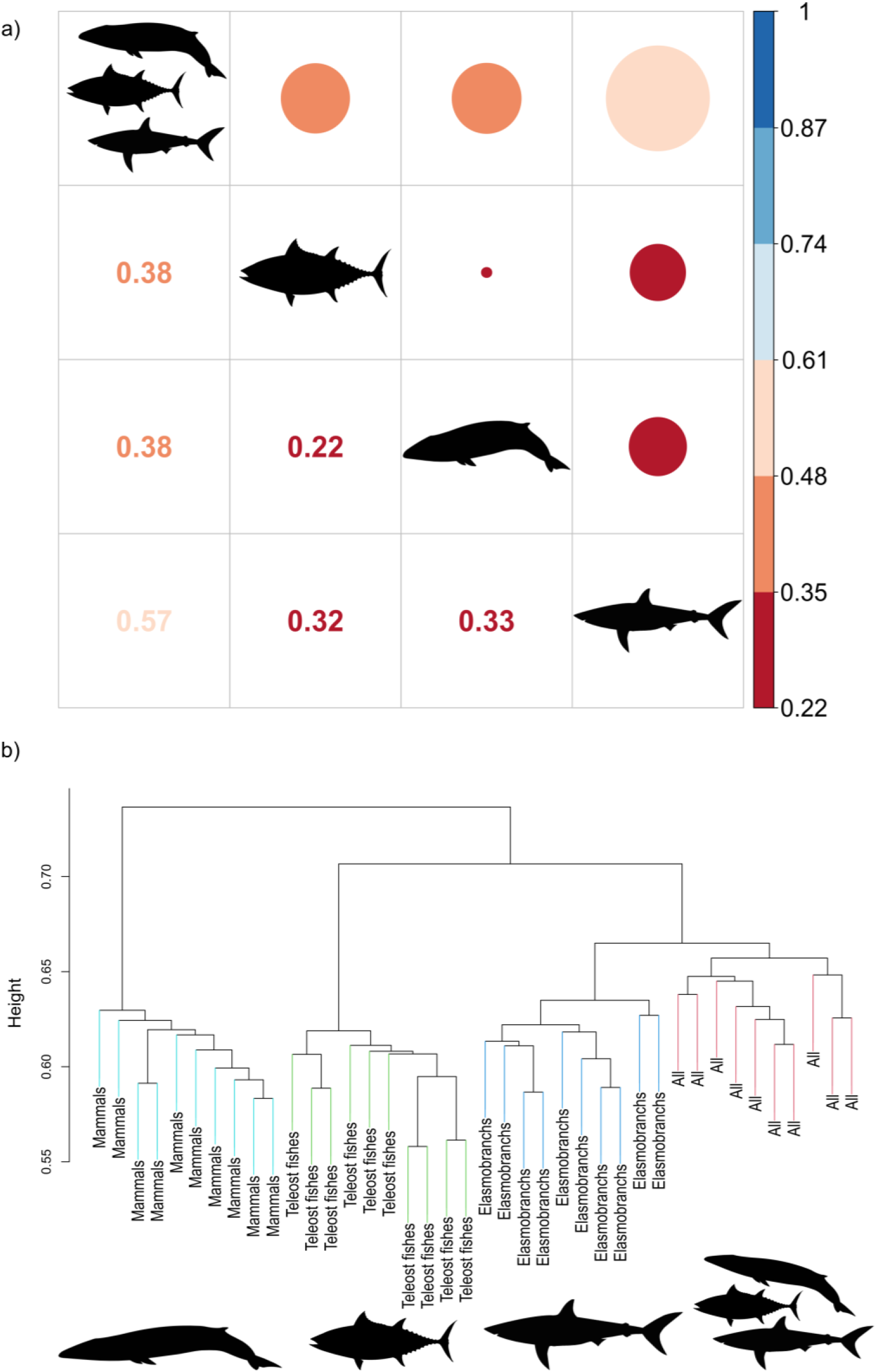
a) Cohen’s Kappa coefficient showing similarity between selection frequency classes across the different conservation feature scenarios. b) Dendrogram displaying the average Jaccard distances between the ten best solutions across the different taxa conservation feature scenarios.

Including all taxa resulted in solutions with highest costs and greatest number of planning units (Supplementary Figure S1). Marine mammal prioritisations required the lowest costs (81,939 ±2206) despite requiring the greatest number of planning units (5644 ±178) whilst elasmobranchs have the greatest costs (104,217 ±2043) despite needing a relatively similar number of planning units to mammals (5526 ±180).

The Mediterranean MPA network does not fulfil the 30 % coverage target for any of the marine predator taxa distributions (Figure 5). Teleost fish distributions overlap most with the current MPA network (24.12 % ±13.27 s.d), whilst marine mammals (16.54 % ±6.49) and elasmobranchs (18.58 % ±10.81) share similar lower levels of protection. Overlap with the MPA network varied greatly for species within a taxonomic group, so the overall differences between groups were not significant (Kruskal-Wallis, p>0.05). Scenarios including a single taxonomic group resulted in greatest overlap with species distributions from that group (Kruskal-Wallis, p < 0.05). Our spatial prioritisation solution for marine mammals afforded the least co-protection to other taxa, with teleost fish distributions overlapping 18.42 % ±11.08 and sharks 19.92 % ±11.06.

**Figure 5.**
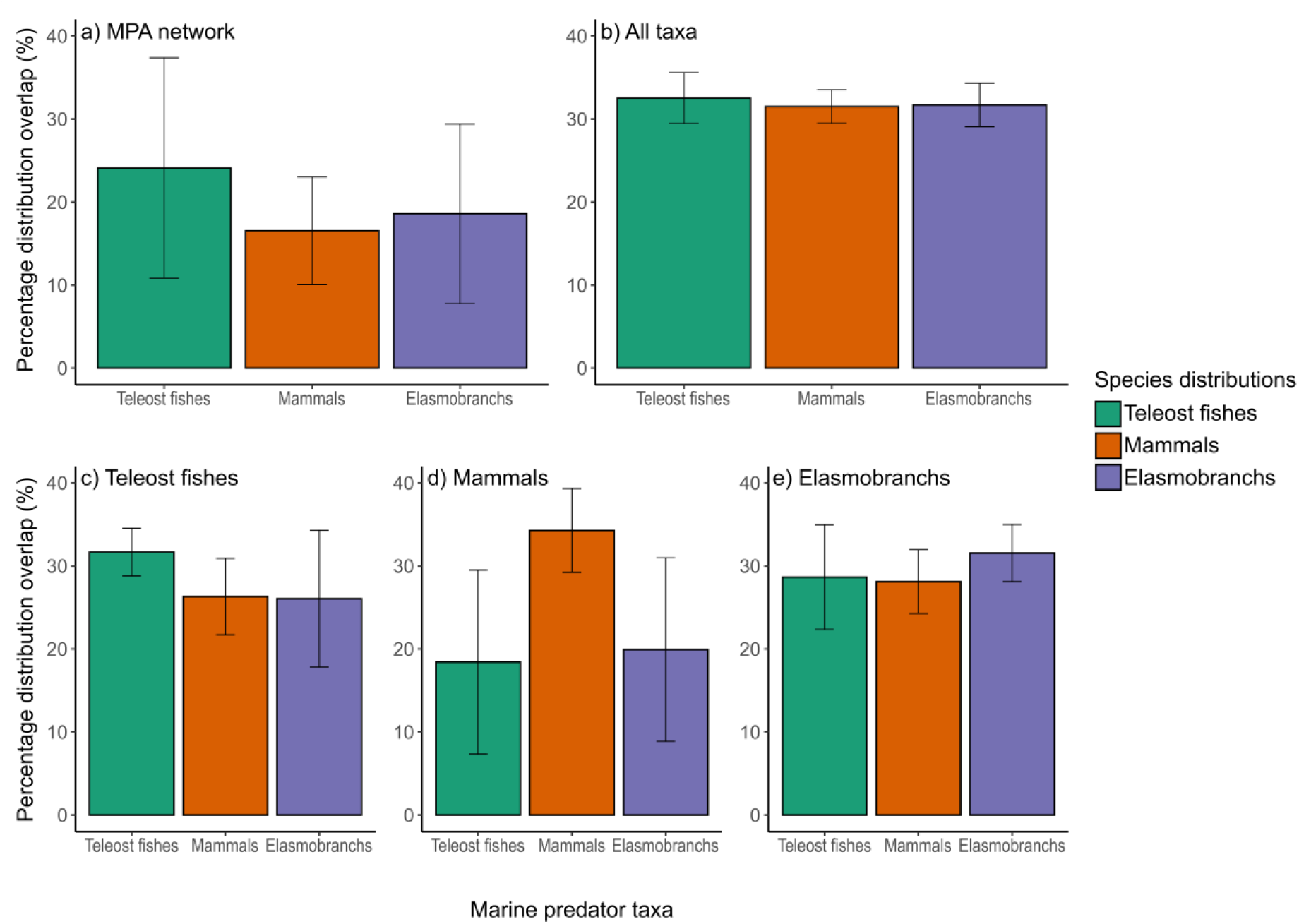
Average percentage distribution overlap of fish, mammal and elasmobranch taxa with a) the current network of Mediterranean MPAs, or conservation feature scenarios with b) all taxa, c) teleost fishes, d) mammals or e) elasmobranchs.

## Discussion

We discover that existing MPA systems in the Mediterranean Sea only afford limited protection to marine predators, with highly variable coverage within and between different taxonomic groups. Instead, marine predator taxa require different conservation priority areas, due to their specific habitat requirements and life histories, as indicated by their heterogeneous distributions. Focussing spatial planning on all species simultaneously, as is common practice, captured the conservation needs well for elasmobranchs, but excluded sites that would gain highest conservation benefits for marine mammals and teleost fishes. Hence, where spatial planning aims to capture all (*i.e*., as many as possible) taxa in conservation management areas, there is a risk of missing conservation needs of important taxa. We advocate that including conservation objectives and actions specific to taxonomic groups will better achieve effective conservation of taxa with contrasting or specialised life histories and habitat needs.

We find a striking contrast between the coastal distributions of sharks and teleost fishes, with the offshore ranges of marine mammals, consistent with globally observed patterns in predator richness (Grady et al., 2019) (Figure 2). These differences may be exaggerated by greater data availability offshore for marine mammals from ferry-based visual surveys, whilst data for elasmobranchs and teleosts largely comes from coastal fisheries (Mannocci et al., 2018; Mancusi et al., 2020). We also find higher predator richness in the north-western basin, due to its proximity to the Strait of Gibraltar which is an important migration corridor connecting the Mediterranean Sea with the Atlantic Ocean (Coll et al., 2010). This result may be inflated by higher observation effort in the north-western basin (Coll et al., 2010). Increasing research efforts to reduce biases in data, and thus ensuring the most effective conservation actions should remain a priority. However, given the threatened statuses and urgent need for conservation measures for marine predators, we must use the best available data and iteratively review and update management as our knowledge improves.

Contrasting distributions of marine predator taxa translated into significant differences in the spatial arrangement of priority areas (Figure 3); highlighting that omitting any of these taxonomic groups from conservation planning will locate MPAs in the wrong areas. Prioritisation solutions including all taxa and elasmobranchs were most similar, driven by elasmobranch species occupying species-poor habitats, or costlier areas which are avoided in the teleost fish or mammal solutions (Kujala et al., 2018). Including representative species across different taxa granted protection to rare species within the taxonomic groups considered.

For example, currently recognised important habitats for critically endangered angel sharks were covered in the elasmobranch priority areas despite the species not being included in the current analysis (Giovos et al., 2022). Most importantly, encompassing all taxa simultaneously provides no information about which species or taxa are covered by which priority areas, making it difficult to implement targeted management measures. The requirement for taxa-specific conservation actions to be incorporated into spatial planning has been acknowledged through ‘Important Marine Mammal Areas’ (IMMAs) and ‘Ecologically and Biologically Significant Marine Areas’ (EBSAs) (Corrigan et al., 2014). However, obligations to act in response to IMMAs/EBSAs are unclear, and it is debatable how they will specifically contribute to area-based conservation (Corrigan et al., 2014). Specific objectives for separate taxa will necessitate less severe restrictions, *i.e*., banning of fishing gear which affects the target taxon, instead of a complete fishing ban (Tixier et al., 2021).

Current Mediterranean MPAs fail to achieve the 30 % coverage target for any of the marine predator taxa (Figure 5). Teleost fish distributions overlap most with Mediterranean MPAs, despite the two largest MPAs, ‘the Pelagos Sanctuary for Marine Mammals’ and ‘Corredor de Migración de Cetaceos del Mediterraneo’, being designated for cetaceans (MedPAN and SPA/RAC, 2022) (Figure 1B). Instead, marine mammal distributions overlapped the least with Mediterranean MPAs, showing misalignment between conservation objectives and outcomes. Currently, the majority of MPAs in the Mediterranean Sea are within European Union waters (Claudet et al., 2020), but our prioritisation solutions highlight important areas for marine predators in the southern Mediterranean Sea, including the Alboran Sea, and along the Tunisian and Egyptian coastlines (Figure 3). The Alboran Sea has been classified as an IMMA with important habitat for threatened cetaceans and overall high cetacean diversity (IUCN-MMPATF). The Tunisian and Egyptian coastlines are both included in EBSAs, and encompass feeding and spawning grounds for fin whales (*Balaenoptera physalus*) and bluefin tuna (*Thunnus thynnus*) respectively (UNEP/CBD/EBSA/WS/2014/3/4, 2014). Since priority areas are not shared equally across countries, cross-country collaborations are required to support those with the highest burden, which will be challenging in the dynamic political environment of the Mediterranean (Mazor et al., 2013). However, this is the most cost-effective method to prioritise key habitats for marine predators, and will improve the likelihood of successful compliance given that stakeholders and conservation features have been considered synergistically (Mazor et al., 2013).

Despite the rapid expansion in the global extent of MPAs, marine predator distributions are not being sufficiently protected, leading to increasing proportions of threatened species. They are notoriously difficult to conserve through MPAs as their vast distributions cannot be encompassed completely, resulting in debate over which key habitats or life history stages should be prioritised in MPA systems. Here, we advocate spatial planning considering marine predator taxonomic groups separately to incorporate their heterogenous distributions arising from divergent life history strategies and habitat use. This approach allows key priority areas for individual taxa to be identified, which are otherwise excluded when considering all taxa concurrently. Designating taxa specific MPAs will enhance the development of more guided management actions to meet the requirements of different taxonomic groups.

## Acknowledgements and data availability

EB was supported by a Leeds Doctoral Scholarship from the University of Leeds. We thank Fabrizio Serena & Monica Barone for providing access to the Mediterranean Large Elasmobranchs Monitoring (Medlem) database. We also greatly appreciate Elena Valsecchi and Antonella Arcangeli for offering feedback on the manuscript.

The authors declare no conflict of interest.

The species distribution models built for this study are available at https://doi.org/10.5061/dryad.280gb5ms5.

## Author contributions

Maria Beger and Elizabeth Boyse led the conceptualisation and design of the study, with Simon J. Goodman supporting this. Elizabeth Boyse performed the data analysis and wrote the first draft. All authors have developed, edited and approved the final manuscript for publication.

