## Supplementary Information for "Marine predator spatial conservation priorities are taxon-specific"

| | |
|---|---|
| **Supplementary Table S1** | List of marine predator species included in stacked species distribution model. |
| **Supplementary Table S2** | Evaluation metrics for marine predator stacked-species distribution model. |
| **Supplementary Table S3** | Importance of environmental predictors in the marine predator stacked-species distribution model. |
| **Supplementary Figure S1** | Mean cost and mean number of planning units across 10 best Marxan solutions for each of the conservation feature scenarios. |
| **Supplementary Figure S2** | Bathymetric map of the Mediterranean Sea displaying Ecologically or Biological Significant areas (EBSAs) and Important Marine Mammal Areas (IMMAs). |
| **Supplementary Table S4** | Kruskal Wallis tests comparing overlap between MPAs or conservation feature scenarios targeting mammals, sharks or fishes and marine predator taxa distributions. |

**Supplementary Table S1**

Species included in stacked species distribution models, displaying the taxonomic group, total length (cm), trophic level and number of occurrences used in the model.

| Species | Common name | Taxa | Total length (cm) | Trophic Level | Number of occurrences |
|---|---|---|---|---|---|
| *Conger conger* | European Conger | Fishes | 573.3 | 4.3 | 649 |
| *Dentex dentex* | Common dentex | Fishes | 100 | 4.5 | 372 |
| *Echelus myrus* | Painted eel | Fishes | 100 | 4.3 | 45 |
| *Epinephelus aeneus* | White grouper | Fishes | 120 | 4 | 49 |
| *Epinephelus marginatus* | Dusky grouper | Fishes | 150 | 4.4 | 969 |
| *Fistularia commersonii* | Bluespotted cornetfish | Fishes | 160 | 4.3 | 63 |
| *Lophius piscatorius* | Angler | Fishes | 200 (SL) | 4.5 | 602 |
| *Merluccius merluccius* | European hake | Fishes | 140 | 4.4 | 1179 |
| *Mola mola* | Ocean sunfish | Fishes | 333 | 3.3 | 3530 |
| *Molva dypterygia* | Blue ling | Fishes | 155 | 4.5 | 119 |
| *Muraena helena* | Mediterranean moray | Fishes | 150 | 4.2 | 1061 |
| *Ophisurus serpens* | Serpent eel | Fishes | 250 | 4.1 | 47 |
| *Pomatomus saltatrix* | Bluefish | Fishes | 130 | 4.5 | 82 |
| *Seriola dumerili* | Greater amberjack | Fishes | 190 | 4.5 | 118 |
| *Sphyraena sphyraena* | European barracuda | Fishes | 165 | 4 | 159 |
| *Sphyraena viridensis* | Yellowmouth barracuda | Fishes | 128 (FL) | 4.3 | 54 |
| *Thunnus alalunga* | Albacore | Fishes | 140 (FL) | 4.3 | 270 |
| *Thunnus thynnus* | Bluefin tuna | Fishes | 458 | 4.5 | 177 |
| *Xiphias gladius* | Swordfish | Fishes | 455 | 4.5 | 48 |
| *Zu cristatus* | Scalloped ribbonfish | Fishes | 118 (SL) | 4.5 | 204 |
| *Alopias vulpinas* | Common thresher | Elasmobranchs | 573.3 | 4.5 | 114 |
| *Carcharhinus longimanus* | Oceanic whitetip shark | Elasmobranchs | 400 | 4.2 | 77 |
| *Cetorhinus maximus** | Basking shark | Elasmobranchs | 1520 | 3.2 | 142 |
| *Dasyatis pastinaca** | Common stingray | Elasmobranchs | 64 (WD) | 4.1 | 163 |
| *Echinorhinus brucus* | Bramble shark | Elasmobranchs | 310 | 4.4 | 41 |
| *Hexanchus griseus* | Bluntnose sixgill shark | Elasmobranchs | 482 | 4.5 | 140 |
| *Isurus oxyrinchus* | Short-fin mako shark | Elasmobranchs | 445 | 4.5 | 81 |
| *Mobula mobular** | Giant devil ray | Elasmobranchs | 520 (WD) | 3.7 | 874 |

| | | | | | |
|---|---|---|---|---|---|
| *Myliobatis aquila** | Common eagle ray | Elasmobranchs | 183 (WD) | 3.6 | 119 |
| *Prionace glauca* | Blue shark | Elasmobranchs | 400 | 4.4 | 324 |
| *Raja clavata** | Thornback ray | Elasmobranchs | 139 | 3.8 | 431 |
| *Squalus acanthias** | Spiny dogfish | Elasmobranchs | 95 | 4.4 | 78 |
| *Torpedo marmorata* | Marbled electric ray | Elasmobranchs | 100 | 4.5 | 116 |
| *Balaenoptera physalus* | Fin whale | Mammals | 2700 | 3.2-4.3 | 1245 |
| *Delphinus delphis* | Short-beaked common dolphin | Mammals | 260 | 4.5 | 1693 |
| *Globicephala melas* | Long-finned pilot whale | Mammals | 670 | 4.5 | 1147 |
| *Grampus griseus* | Risso’s dolphin | Mammals | 380 | 4.36-4.54 | 410 |
| *Orcinus orca* | Killer whale | Mammals | 980 | 4.5-4.6 | 115 |
| *Physeter macrocephalus* | Sperm whale | Mammals | 2400 | 4.5-4.7 | 2307 |
| *Stenella coeruleoalba* | Striped dolphin | Mammals | 260 | 4.5 | 7822 |
| *Tursiops truncatus* | Bottlenose dolphin | Mammals | 380 | 4.5 | 3991 |
| *Ziphius cavirostris* | Cuvier’s beaked whale | Mammals | 750 | 4.5 | 113 |

**Supplementary Table S2**

Pearson correlation coefficient calculated from the difference between a full model and one with each environmental variable omitted in turn for individual species models then averaged across species.

| | Mean bathymetry | Mean sea surface temperature | Mean chlorophyll concentration | Mean temperature range | Bathymetric slope | Distance from shore |
|---|---|---|---|---|---|---|
| Mean | 14.59 | 31.76 | 13.17 | 16.7 | 8.31 | 15.47 |
| SD | 6.56 | 13.74 | 5.1 | 5.53 | 1.96 | 8.32 |

**Supplementary Table S3**

Results from six metrics used to evaluate the prediction accuracy of species assemblage predictions in the ‘perfect knowledge’ SSDM.

| | Species Richness Error | Prediction Success | Kappa | Specificity | Sensitivity | Jaccard |
|---|---|---|---|---|---|---|
| Mean | 19.06 | 5.98 | 0.995029 | 0.544447 | 0.983425 | 0.065596 |
| SD | 7.229589 | 1.797052 | 0.000441 | 0.172317 | 0.124641 | 0.059542 |

**Supplementary Figure S1**

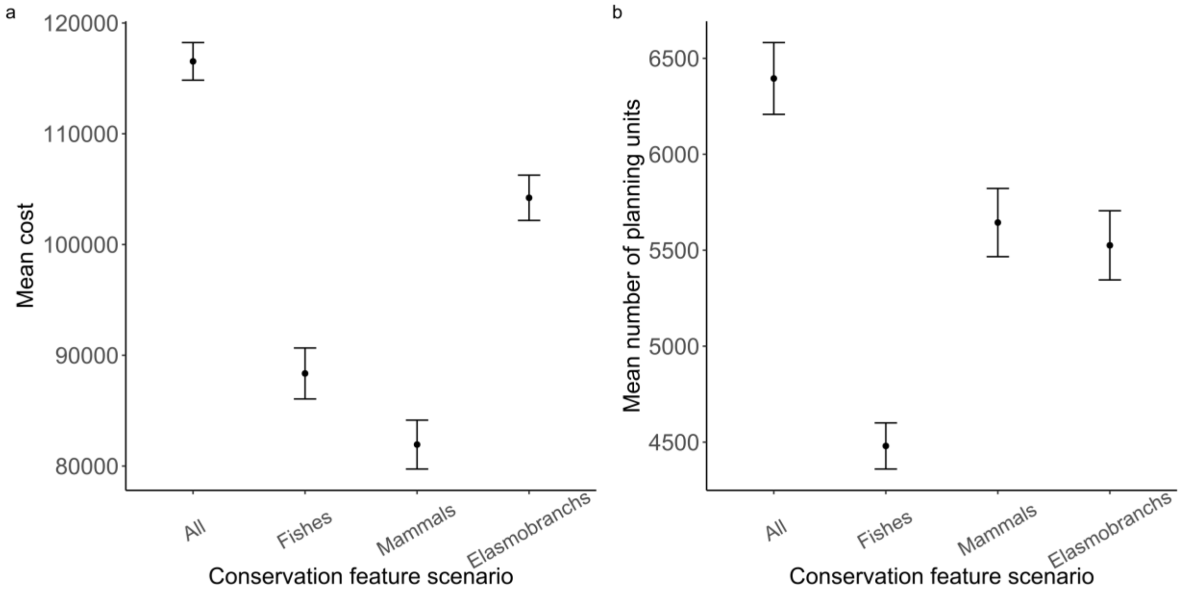


a) Mean cost across 10 best Marxan solutions for each of the conservation feature scenarios. b) Mean number of planning units in solutions from 10 best Marxan runs for each of the conservation feature scenarios.

**Supplementary Figure S2**

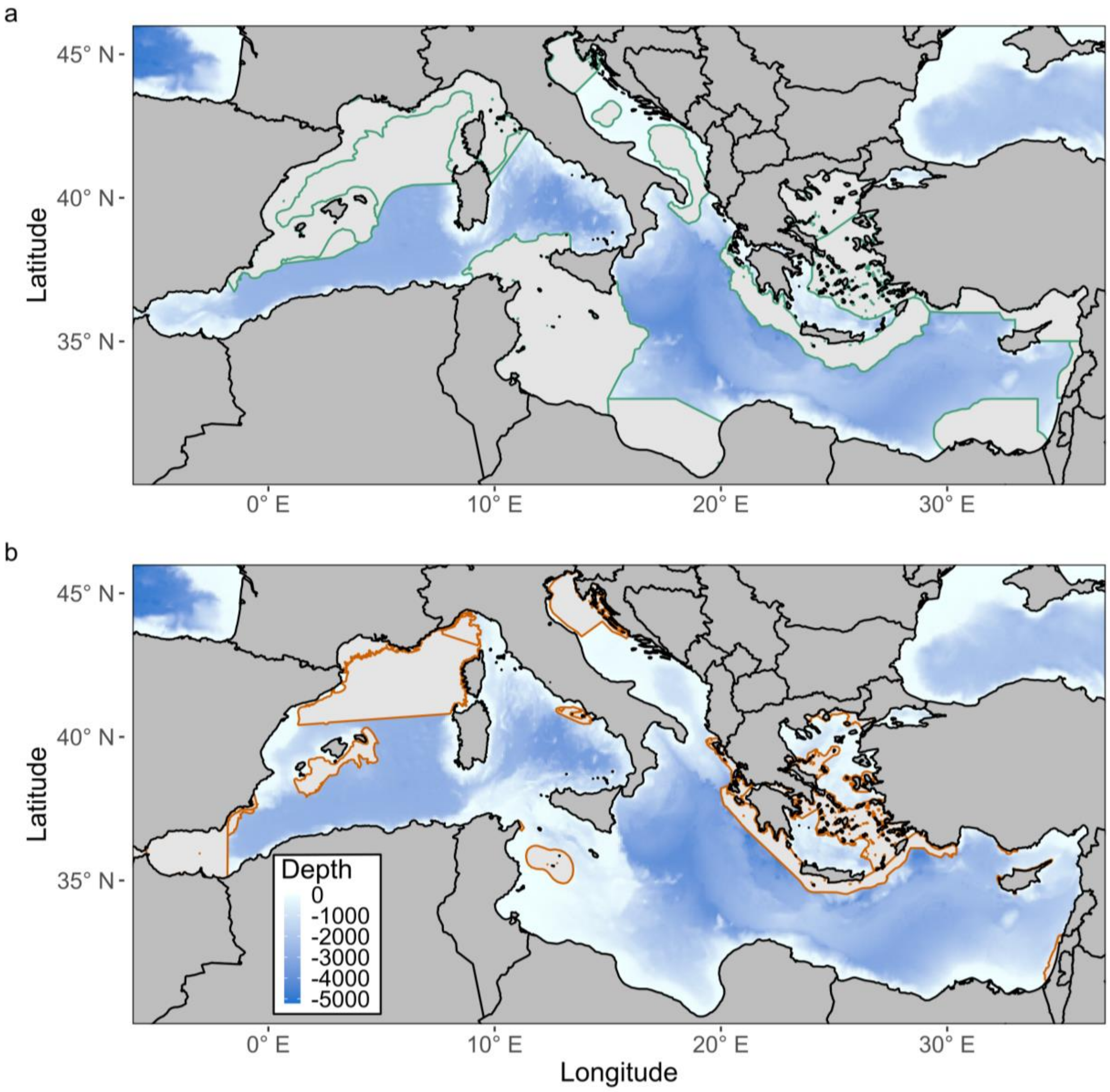


Bathymetric map of the Mediterranean Sea displaying a) Ecologically or Biological Significant areas (EBSAs) and b) Important Marine Mammal Areas (IMMAs).

**Supplementary Table S4**

Kruskal Wallis to compare the overlap between MPAs or conservation feature scenarios for mammals, fishes and sharks and species distributions of individual taxa (mammals, fishes, sharks).

| | Chi-squared | Degrees of freedom | P value |
|---|---|---|---|
| MPAs | 3.2742 | 2 | 0.1945 |
| Mammals | 108.43 | 2 | <0.001* |
| Fishes | 142.49 | 2 | <0.001* |
| Sharks | 55.629 | 2 | <0.001* |
